# Fluid flow sensitizes bacterial pathogens to chemical stress

**DOI:** 10.1101/2022.09.07.506966

**Authors:** Gilberto C. Padron, Alexander M. Shuppara, Anuradha Sharma, Matthias D. Koch, Jessica-Jae S. Palalay, Jana N. Radin, Thomas E. Kehl-Fie, James A. Imlay, Joseph E. Sanfilippo

## Abstract

Cells regularly experience fluid flow in natural systems. However, most experimental systems rely on batch cell culture and fail to consider the effect of flow-driven dynamics on cell physiology. Using microfluidics and single-cell imaging, we discover that the interplay of physical shear rate (a measure of fluid flow) and chemical stress trigger a transcriptional response in the human pathogen *Pseudomonas aeruginosa*. In batch cell culture, cells protect themselves by quickly scavenging the ubiquitous chemical stressor hydrogen peroxide (H_2_O_2_) from the media. In microfluidic conditions, we observe that cell scavenging generates spatial gradients of H_2_O_2_. High shear rates replenish H_2_O_2_, abolish gradients, and generate a stress response. Combining mathematical simulations and biophysical experiments, we find that cells in flow are sensitive to a H_2_O_2_ concentration that is 100-1000 times lower than traditionally studied in batch cell culture. Surprisingly, the shear rate and H_2_O_2_ concentration required to trigger a transcriptional response closely match their respective values in the human bloodstream. Thus, our results explain a long-standing discrepancy between H_2_O_2_ levels in experimental and natural systems. Finally, we demonstrate that the shear rate and H_2_O_2_ concentration found in the human bloodstream trigger gene expression in the blood-relevant human pathogen *Staphylococcus aureus*, suggesting that flow sensitizes bacteria to chemical stress in natural environments.

## Main Text

Classically, research on bacterial stress responses has focused on chemical stressors such as nutrient availability^1^, pH^2^, antibiotics^3^, and oxidative stress^4^. Bacteria respond to chemical perturbations with well-studied physiological responses to survive and grow in stressful situations^5–8^. For simplicity, bacterial stress responses have been largely studied in batch cell culture, which has allowed researchers to identify and characterize many of the important signaling pathways associated with stress. However, simplified experimental systems neglect the dynamic mechanical features of natural and host systems^9^. Recently, a surge of research on bacterial mechanosensing has revealed that fluid flow impacts virulence^10,11^, biofilm formation^12^, and gene expression^13^. While bacterial transcriptional responses to flow were assumed to require the measurement of forces^10,12^, one report challenged this assumption and revealed that flow can trigger bacterial gene expression in a force-independent manner^13^. This report proposed naming flow-sensitive transcriptional responses “rheosensitive” (as *rheo*- is Greek for flow), due to lack of direct evidence that bacteria respond to flow by measuring forces^13^. Thus, it is currently debated how flow generates bacterial transcriptional responses.

To understand how flow affects gene expression, we focused on the flow-sensitive transcriptional response in the bacterium *Pseudomonas aeruginosa. P. aeruginosa* responds to flow by upregulating a suite of genes^13^, including many genes upregulated during human infection^14^. As a representative example of the larger flow-sensitive response, we focused our efforts on the *fro* operon. The *fro* operon is rapidly and robustly upregulated by flow^13^ and required for full infection in multiple animal models^15,16^. For single-cell analysis, we used a *fro* reporter strain that reports on *fro* expression with yellow fluorescent protein (YFP) and encodes a constitutively expressed mCherry for normalization. To confirm that flow induces *fro* expression, cells in a microfluidic device were simultaneously subjected to flow from a syringe pump and imaged with a fluorescence microscope (Figure 1A). Throughout this study, we represent the intensity of flow using shear rate, which is calculated using flow rate and channel dimensions^13^. Consistent with previous results^13^, *fro* is strongly induced after 3 hours of exposure to a shear rate of 800 sec^−1^ in LB medium (Figure 1B). Thus, the *fro* reporter represents a valuable tool to dissect how flow generates transcriptional responses in bacteria.

**Figure 1:**
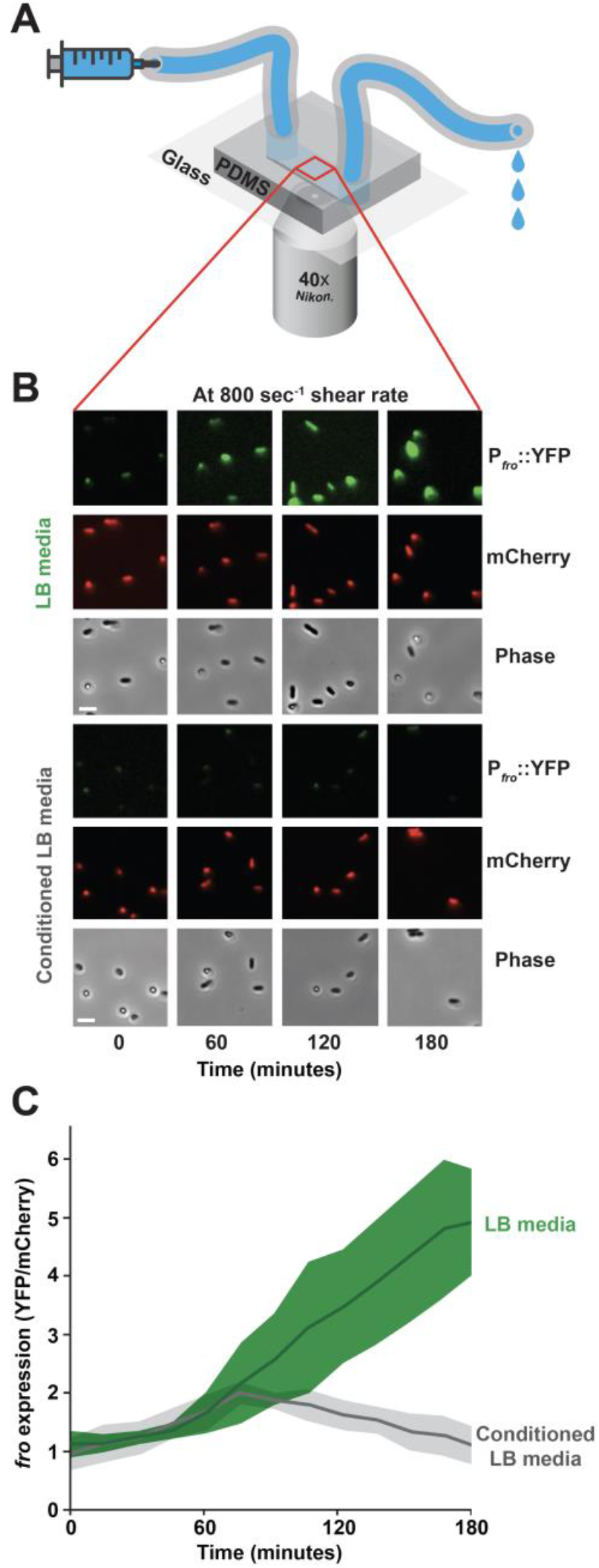
Conditioning media suppresses flow-sensitive gene expression in *Pseudomonas aeruginosa*. **(A)** Microfluidic setup used in this study. Microfluidic devices are custom fabricated with polydimethylsiloxane (PDMS) and glass coverslips. Channels are 50 μm tall x 500 μm wide. Bacteria adhere to the channel wall. Microfluidic devices are simultaneously subjected to flow from a syringe pump and imaged by a fluorescence microscope. (**B)** Fluorescence and phase images of the *P. aeruginosa fro* reporter strain in flow (at a shear rate of 800 s^−1^) over 180 minutes. Images are representative of three biological replicates. Scale bars, 5 μm. **(C)** Quantification of *fro* expression by dividing YFP intensity by mCherry intensity as described in Figure S1. Green represents cells exposed to LB media, while gray represents cells exposed to conditioned LB. Shaded regions show standard deviation of three biological replicates.

How does flow generate a transcriptional response? One possibility is that flow affects a rate-dependent biophysical process such as chemical transport. To test this hypothesis, we flowed fresh LB medium and LB medium that had been conditioned by *P. aeruginosa* into independent channels of the device. Conditioned medium was generated by exposing fresh, sterile LB to a culture of *P. aeruginosa* for 60 minutes, followed by filter-sterilizing the media to remove cells. To quantify reporter intensity during flow treatment, we optimized and employed a MATLAB-based program^17^ to identify cells and quantify single cell fluorescence intensity (Figure S1). We determined that cells exposed to flow with LB induced *fro* expression 5-fold, but *fro* was not induced after 3 hours of exposure to flow in conditioned LB (Figure 1B, 1C). Thus, our results suggest that chemical transport of a molecule into or out of cells underlies how bacteria respond to flow.

What is the identity of the flow-sensitive molecule? To examine if the molecule was conserved, we tested the effect of conditioning media with *Escherichia coli, Staphylococcus aureus*, and *Enterococcus faecalis*. Conditioning media with *E. coli* or *S. aureus* led to complete loss of *fro* induction in flow, while conditioning media with *E. faecalis* resulted in high *fro* induction (Figure 2A). *P. aeruginosa, E. coli*, and *S. aureus* are catalase-positive^18^, while *E. faecalis* is catalase-negative^19^. Catalase specifically degrades H_2_O_2_^20^, supporting the hypothesis that the flow-sensitive molecule is H_2_O_2_. We reasoned that if H_2_O_2_ was the flow-sensitive molecule, it must be present in our media. We tested H_2_O_2_ levels with a peroxidase assay^21^ and determined that our laboratory LB stocks contained approximately 9 μM H_2_O_2_ (Figure S2). The presence of H_2_O_2_ in LB is a reproducible but underappreciated detail^22,23^. The chemical production of H_2_O_2_ in LB is mediated by a light-dependent reaction involving riboflavin^23^. Consistent with the literature^18,19^, we confirm that *P. aeruginosa, E. coli*, and *S. aureus* can deplete H_2_O_2_ from media in 30 minutes, while *E. faecalis* cannot (Figure S3). To directly test the role of catalase, we treated LB with purified catalase, which depleted H_2_O_2_ concentrations in LB to effectively zero (Figure S4). Catalase-treated media did not induce *fro* expression in flow, further supporting the hypothesis that the flow-sensitive molecule is H_2_O_2_. (Figure 2B). Finally, we tested *fro* expression with conditioned media re-supplied with 9 μM H_2_O_2_. Re-introduction of H_2_O_2_ restored robust *fro* induction (Figure 2B), demonstrating that the flow-sensitive molecule is H_2_O_2_.

**Figure 2:**
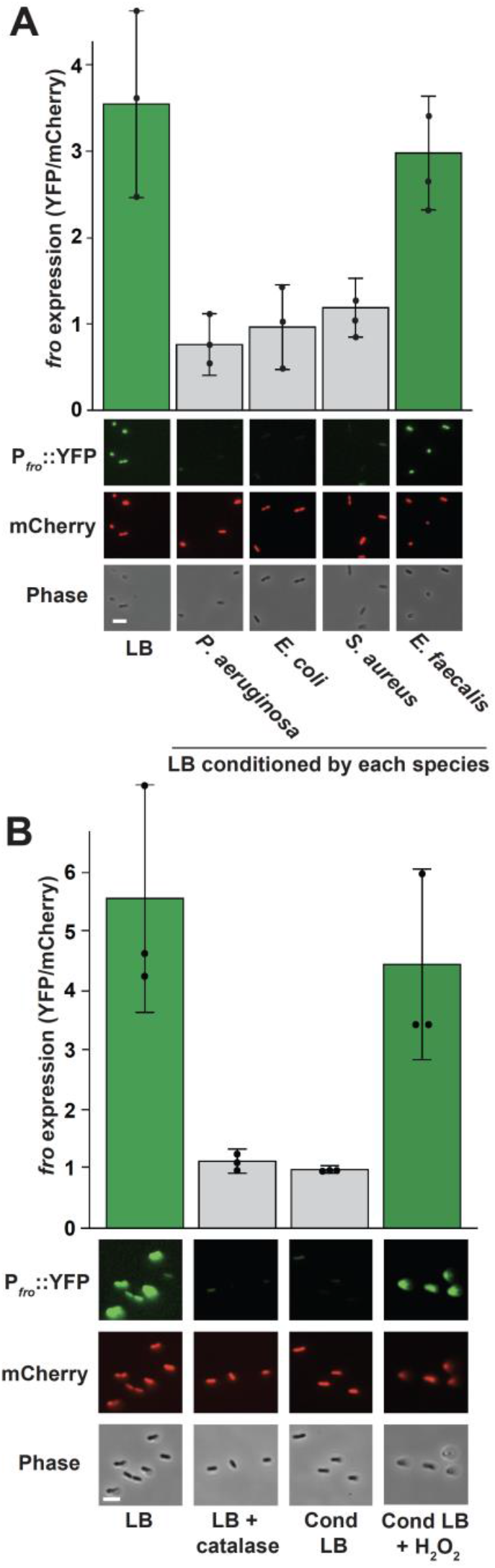
H_2_O_2_ is a flow-sensitive molecule. **(A)** *P. aeruginosa fro* expression after 180 minutes in flow (at a shear rate of 800 s^−1^) with LB media, or media conditioned by *P. aeruginosa, E. coli, S. aureus*, or *E. faecalis* for 60 minutes. Quantification shows the average and standard deviation of three biological replicates. Images show YFP and mCherry fluorescence, as well as phase contrast. **(B)** *fro* expression after 180 minutes in flow (at a shear rate of 800 s^−1^) with LB media, LB treated with catalase, media conditioned by *P. aeruginosa* for 60 minutes, and conditioned media resupplied with 9 μM H_2_O_2_. Quantification shows the average and standard deviation of three biological replicates. Images show YFP and mCherry fluorescence, as well as phase contrast. Scale bars, 5 μm. Channels are 50 μm tall x 500 μm wide.

Armed with our understanding that H_2_O_2_ is the flow-sensitive molecule, we re-analyzed RNA-sequencing data from a previous microfluidic-based transcriptomic experiment^13^. In addition to the *fro* operon, we identified many flow-induced genes implicated in H_2_O_2_ scavenging, such as *ahpCF, ahpB*, and *katB* (Figure 3A). AhpCF and AhpB are NADH peroxidases that scavenge H_2_O_2_^24–26^. KatB is a catalase that converts H_2_O_2_ into water and oxygen^20,24^. AhpCF, AhpB, and KatB are induced by H_2_O_2_ and regulated by OxyR, a H_2_O_2_ sensor located in the cytoplasm^24,27–30^. Thus, our results suggest that shear rate leads to the intracellular accumulation of H_2_O_2_. In support of this model, an additional H_2_O_2_-regulated gene *trxB2*, which encodes thioredoxin reductase^27^, was also upregulated in response to shear rate (Figure 3A). Together, our results suggest the interplay of shear rate and H_2_O_2_ generate cellular stress and trigger a transcriptional response.

**Figure 3:**
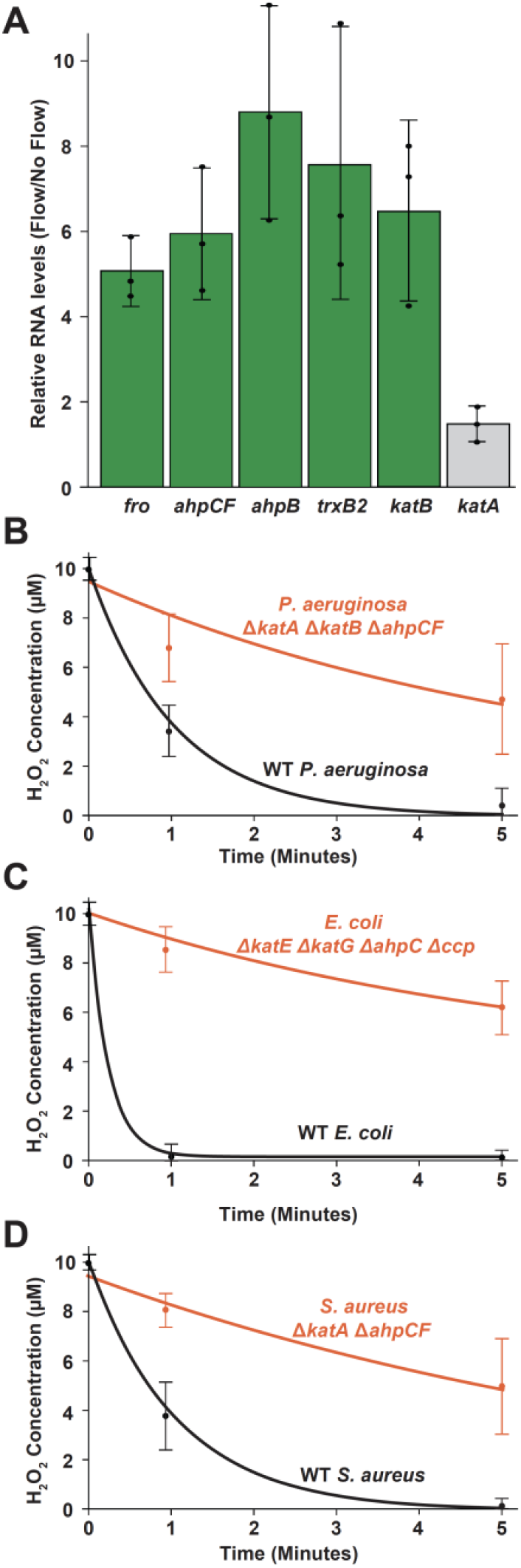
Fluid flow upregulates multiple enzymes that scavenge H_2_O_2_. **(A)** *P. aeruginosa* RNA sequencing data^13^ showing relative transcript levels comparing flow conditions to no flow conditions. *fro* represents an average of the *froABCD* operon and *ahpCF* represents an average of *ahpC* and *ahpF*. Quantification shows the average and standard deviation of three biological replicates. **(B)** H_2_O_2_ concentration of LB media over time when treated with wild-type *P. aeruginosa* and Δ*katA*Δ*katB*Δ*ahpCF* mutant cells at an OD of 0.4-0.5. **(C)** H_2_O_2_ concentration of LB media over time when treated with wild-type *E. coli* and Δ*katE*Δ*katG*Δ*ahpC*Δ*ccp* mutant cells at an OD of 0.4-0.5. **(D)** H_2_O_2_ concentration of LB media over time when treated with wild-type *S. aureus* and Δ*katA*Δ*ahpCF* mutant cells at an OD of 0.4-0.5. All H_2_O_2_ concentrations were measured using a peroxidase assay^21^ and quantification shows the average and standard deviation of three biological replicates.

What is the relationship between flow and H_2_O_2_? We hypothesize that shear rate replenishes H_2_O_2_ to overcome rapid scavenging by cells. To explore the kinetics of H_2_O_2_ scavenging, we used a peroxidase assay^21^ to measure H_2_O_2_ concentrations of LB media treated with *P. aeruginosa, E. coli*, or *S. aureus* over 60 minutes. Additionally, we compared wild-type and mutant strains of *P. aeruginosa, E. coli*, and *S. aureus*. At a cell density of 0.5 OD, wild-type*P*. *aeruginosa* scavenged about 50% of the available H_2_O_2_ in 30 seconds, while a *ΔkatA* Δ*katB* Δ*ahpCF* mutant was significantly impaired at scavenging H_2_O_2_ (Figure 3B). Wild-type *E. coli* scavenges about 90% of the available H_2_O_2_ in 30 seconds, while an Δ*ahpC* Δ*katG* Δ*katE* Δ*ccp* mutant had significantly diminished scavenging ability^31^ (Figure 3C). Finally, wild-type *S. aureus* scavenged about 25% of the available H_2_O_2_ in 30 seconds and a *ΔkatA* Δ*ahpCF* mutant was also significantly impaired at H_2_O_2_ scavenging (Figure 3D). Together, our batch cell culture experiments reveal that bacteria rapidly scavenge H_2_O_2_, providing support to our hypothesis that flow triggers a biological response by replenishing H_2_O_2_.

How sensitive are bacterial cells to H_2_O_2_ in flow? We measured *fro* induction at a range of H_2_O_2_ concentrations, while maintaining a constant shear rate of 800 sec^−1^. We observed that 2 μM H_2_O_2_ did not induce *fro* expression. In contrast, 4 μM H_2_O_2_ led to a 3-fold induction and 8 μM H_2_O_2_ led to a 5-fold induction in *fro* expression (Figures 4A, S5A). Thus, the minimum concentration necessary to elicit a flow-sensitive response at a shear rate of 800 sec^−1^ is approximately 4 μM H_2_O_2_ (Figure 4A). Surprisingly, 4 μM is 100-1000 times lower than the H_2_O_2_ concentration traditionally studied in batch cell culture^24,32–36^. Moreover, the H_2_O_2_ concentration in the human bloodstream is thought to be between 1 and 5 μM^37^. Together, these results support the hypothesis that bacteria in flow are highly sensitive to H_2_O_2_ and suggests that flow generates stress in natural environments.

**Figure 4:**
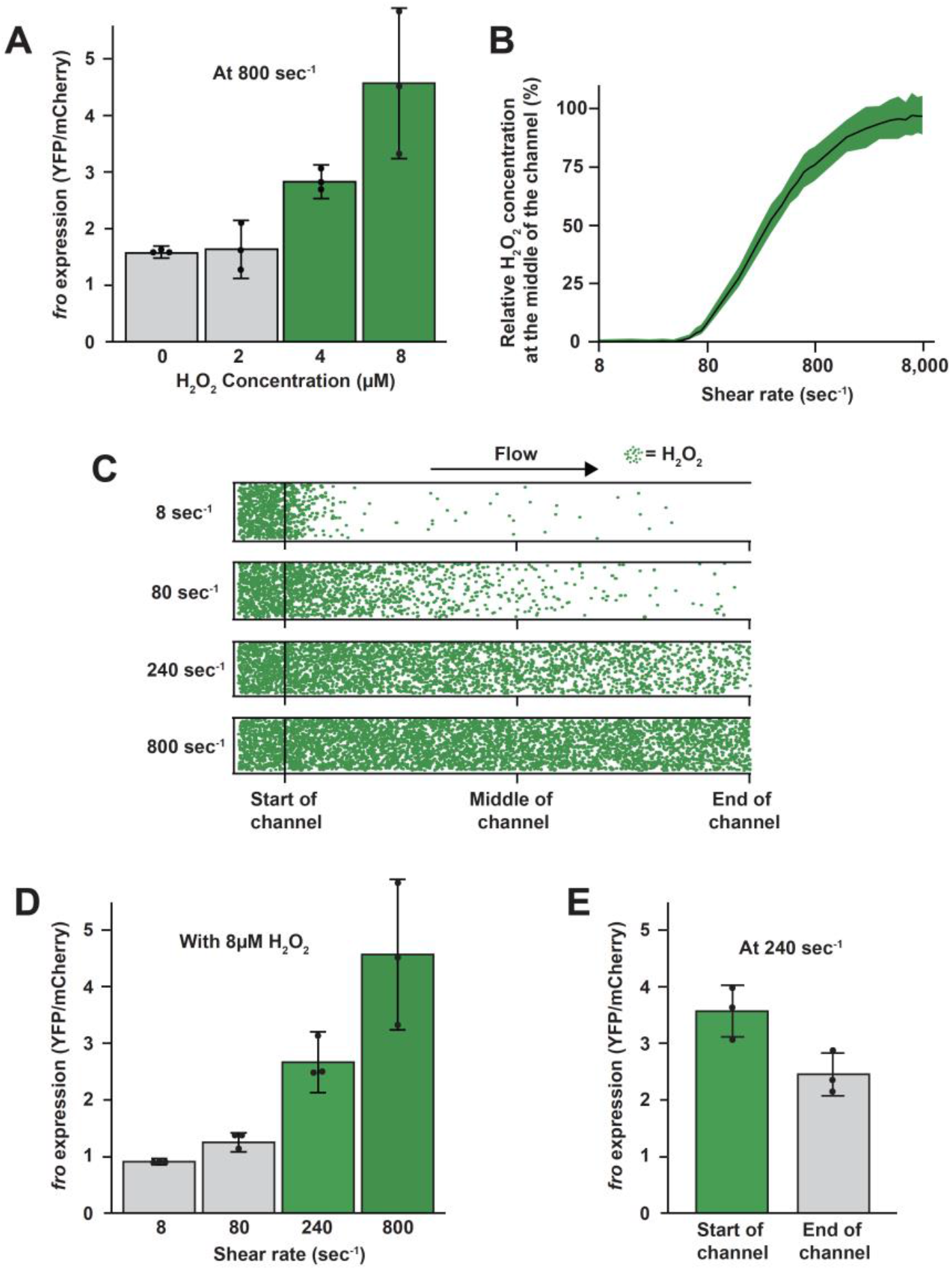
Simulation and experiment determine the minimal shear rate to trigger flow-sensitive gene expression. **(A)** *P. aeruginosa fro* expression after 180 minutes in flow (at a shear rate of 800 s^−1^) with LB media with varied H_2_O_2_ concentrations. **(B)** Relationship between shear rate and the relative H_2_O_2_ concentrations in the middle of a channel from our simulations. Quantification shows the average and standard deviation of multiple bins in the middle of the channel. **(C)** Visual representation of H_2_O_2_ molecules in simulated microfluidic channels at different shear rates. Green dots represent individual H_2_O_2_ molecules experiencing diffusion and fluid flow from left to right. To represent bacterial scavenging, 1 of every 100 molecules that hit the lower surface is removed. This assumption is based on calculations that use the scavenging kinetics defined in Figure 3. **(D)** *fro* expression after 180 minutes in flow (at 8 μM H_2_O_2_) with varied shear rates. **(E)** *fro* expression at the start and end of a 27cm long channel after 180 minutes in flow at 8 μM H_2_O_2_ with a shear rate of 240 s^−1^. Quantification in panels A, D, E shows the average and standard deviation of three biological replicates. Green and gray in panels A, D, E signify expression levels that are statistically different with P<0.05.

To understand the mathematical relationship between flow and H_2_O_2,_ we calculated the Péclet number for our experimental system. The Péclet number describes if shear rate or diffusion is dominant in a particular regime and is proportional to the shear rate divided by diffusion. When shear rate dominates diffusion, the Péclet number is greater than 1. When diffusion dominates shear rate, the Péclet number is less than 1. To solve for the minimal shear rate required to overtake diffusion, we set the Péclet equation equal to 1 and solved for shear rate (Figure S6). Our calculations show that a shear rate of approximately 166 sec^−1^ should dominate diffusion in our experiment system (Figure S6).

To generate testable mathematical predictions, we developed a simulation for the advection-diffusion transport of individual H_2_O_2_ molecules in a channel. The simulation uses experimental values for shear rate and the estimated diffusion coefficient of H_2_O_2_^38^. To simulate H_2_O_2_ removal by cells, we included a feature where 1 of every 100 molecules that contacted the channel surface was removed. We used the value of 1 in 100 based on calculations that use the scavenging kinetics defined in Figure 3. Our simulation shows that as shear rate increases from 8 to 8,000 sec^−1^, the H_2_O_2_ concentration at the middle of the channel increases (Figure 4B). For low shear rates, H_2_O_2_ molecules are rapidly depleted. In contrast, higher shear rates replenish depleted molecules and maintain the concentration of H_2_O_2_ close to the initial value. Our simulated results highlight how rapid H_2_O_2_ scavenging generates gradients of H_2_O_2_, which are abolished by shear rates of at least 240 sec^−1^. Together, our mathematical calculations and simulations predict that a shear rate of 166 sec^−1^ to 240 sec^−1^ will trigger flow-sensitive gene expression.

To test our mathematical predictions, we measured *fro* induction at a range of shear rates, while maintaining a constant H_2_O_2_ concentration of 8 μM. We observed that shear rates of 8 sec^−1^ and 80 sec^−1^ did not trigger *fro* expression. In contrast, 240 sec^−1^ led to a 3-fold induction and 800 sec^−1^ led to a 5-fold induction in *fro* expression (Figure 4D, S5B). Thus, the minimum shear rate necessary to elicit a flow-sensitive response at a H_2_O_2_ concentration of 8 μM is approximately 240 sec^−1^ (Figure 4D). For perspective, human veins have shear rates of approximately 80 sec ^−1^ and human arteries have shear rates of approximately 800 sec ^−1 39^. To test the prediction that cell scavenging generates spatial H_2_O_2_ gradients, we examined *fro* expression at the beginning and end of a long microfluidic channel (Figure S7). For this experiment, we used a shear rate of 240 sec^−1^ and a H_2_O_2_ concentration of 8 μM. Cells at the beginning of the channel induced *fro* expression 3.5 fold, while cells at the end of channel experienced lower induction levels (Figure 4E). Together, our biophysical experiments support our mathematical predictions, establish that flow modulates chemical gradients, and demonstrate that physiological levels of flow trigger a transcriptional response in *P. aeruginosa*.

Does flow generate stress in other bacteria? The biological requirements for a flow-sensitive response are a permeable membrane, H_2_O_2_ scavenging, and H_2_O_2_ sensing capacity.

As these three features are widespread in bacteria, we hypothesized that the human pathogen *S. aureus* would also exhibit flow-sensitive gene expression. We focused our efforts on *S. aureus* as it infects the human bloodstream, which contains high shear rates^39^ and low micromolar H_2_O_2_ concentrations^37^. We generated a *S. aureus* YFP fluorescent reporter to the promoter of *ahpCF*. In *S. aureus, ahpCF* expression is induced by H_2_O_2_ through the function of the H_2_O_2_-sensing transcriptional regulator PerR^40,41^ (Figure S8). When we subjected *S. aureus* cells to a shear rate of 800 s^−1^ and a H_2_O_2_ concentration of 8 μM, *ahpCF* was induced 4-fold compared to no flow conditions (Figure 5). Thus, flow triggers gene expression in *S. aureus*, suggesting that the flow-sensitive stress response we have discovered is widely conserved.

**Figure 5:**
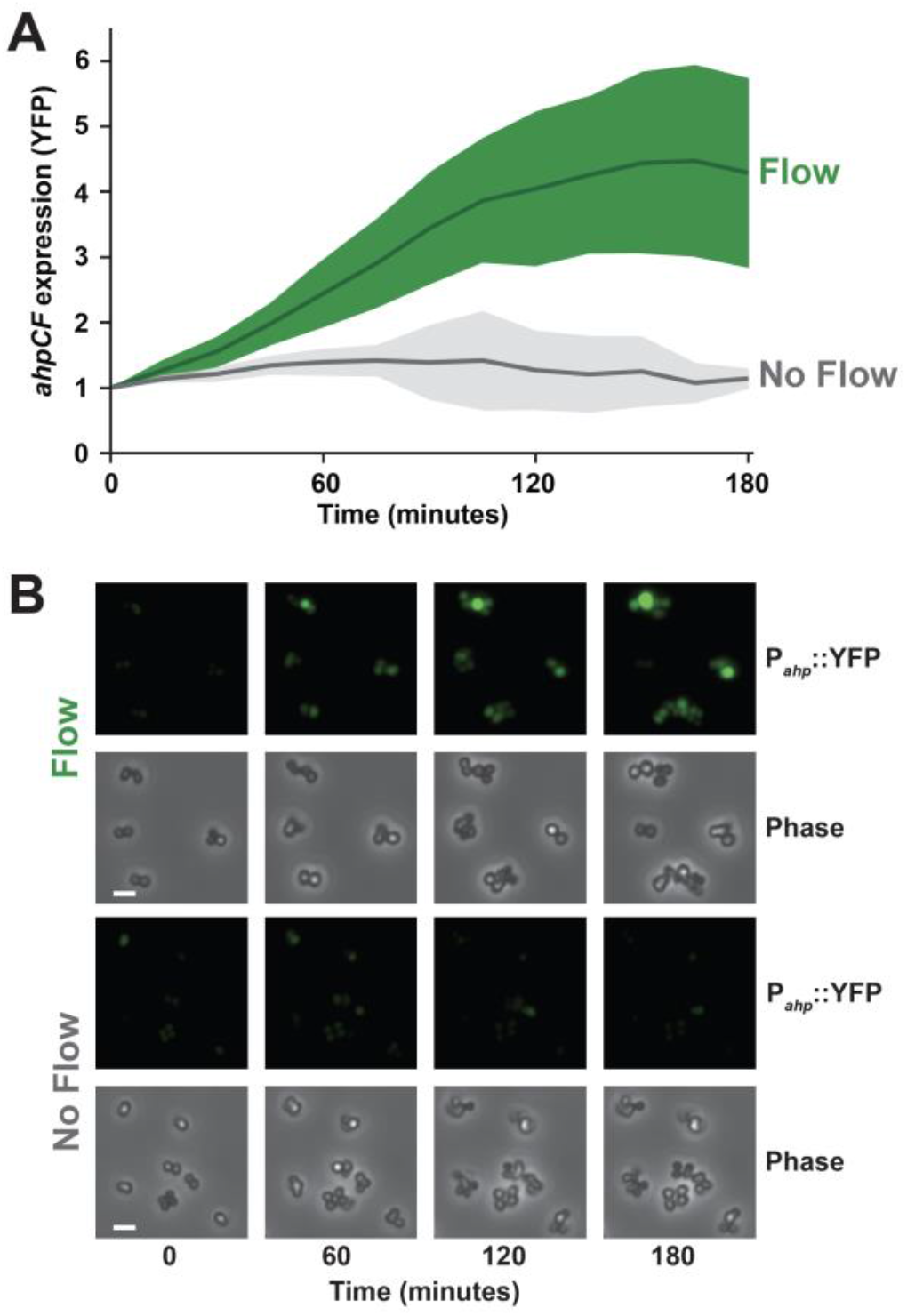
Shear rate triggers flow-sensitive gene expression in *Staphylococcus aureus*. **(A)** *S. aureus ahpCF* expression over 180 minutes in the presence of flow (green) or no flow (gray). Cells contained a P_*ahp*_::YFP transcriptional reporter. Both treatments had 8 μM H_2_O_2_. Flow treatment was at a shear rate of 800 s^−1^. Quantification shows the average and standard deviation of at least three biological replicates. **(B)** YFP fluorescence and phase contrast images representative of at least three biological replicates. Scale bars, 5 μm. Channels are 50 μm tall x 500 μm wide.

Collectively, our results provide a molecular mechanism of how flow generates stress and triggers a transcriptional response (Figure S8). We show that our flow-sensitive transcriptional response hinges on the quantitative relationships between shear rate, H_2_O_2_, and cell scavenging. First, H_2_O_2_ from the environment diffuses into cells, where it is scavenged by catalases and NADH peroxidases (Figure 3). Second, H_2_O_2_ scavenging results in a zone of depletion and H_2_O_2_ spatial gradients in the environment (Figure 4). Third, shear rates above 240 sec^−1^ replenish H_2_O_2_ in the environment, abolish spatial gradients, and lead to increased accumulation of H_2_O_2_ in cells (Figure 4). Fourth, cells sense intracellular H_2_O_2_ levels and activate a transcriptional response that induces factors to mitigate H_2_O_2_ damage (Figure 5). Thus, flow triggers a biological response by countervailing the ability of cells to remove a chemical stressor from the environment.

Until recently^13^, bacterial responses to flow were also assumed to require force-sensing^10,12^. Due to a few well-established examples of force-sensing in eukaryotes^42,43^, the study of bacterial flow responses has focused on the effects of forces and has largely neglected the effects of flow on chemical transport. In contrast, the field of fluid dynamics has long appreciated that flow is a major driver of chemical transport^44^. Our discovery that flow triggers a stress response by replenishing H_2_O_2_ renews the call for caution in the characterization of flow-sensitive responses. Furthermore, our results provide proof-of-principle that flow could amplify or nullify the effect of a wide variety of molecules, such as amino acids, carbon sources, oxygen, and antimicrobials. Thus, flow likely has a far-reaching effect on bacterial physiology that is currently misunderstood and underappreciated.

As an analogy to our findings, we note that flow-sensitive gene expression is conceptually like wind chill. In our system, shear rate generates a biological response by abolishing a chemical gradient. Wind chill describes how wind speed generates a biological response by abolishing a temperature gradient. Historically, the concept of wind chill did not exist until 1945^46^. However, it is now widely reported by the National Weather Service due to the critical importance of its effect on human health. By analogy, we propose that in order to understand how cells interact with their environment in natural systems, it is essential to include flow in experimental systems.

Our discovery that bacteria in flow are sensitive to H_2_O_2_ concentrations that are 100-1000 times lower than traditionally used should provoke a paradigm shift in the way we think about H_2_O_2_ stress. For simplicity, studies on H_2_O_2_ stress in bacteria have primarily used batch cell cultures and millimolar amounts of H_2_O_2_^24,32–36^. However, natural environments are unlikely to contain such high H_2_O_2_ concentrations^9^. Additionally, it has been noted that H_2_O_2_ sensors OxyR and PerR are likely sensitive to much lower concentrations^23,29,45^. Thus, the H_2_O_2_ concentrations required to generate bacterial responses in laboratory conditions do not reflect the environments where bacteria live. Our observation that bacteria in flow are sensitive to low-micromolar levels of H_2_O_2_ likely reconciles this discrepancy. Natural bacterial environments, such as the human bloodstream, contain low-micromolar levels of H_2_O_2_^37^. Without considering the effect of flow, these levels are insufficient to generate bacterial responses. In light of our study, it is apparent that H_2_O_2_ in flowing blood is sufficient to affect bacteria and potentially has a role in the restriction of bacterial growth.

## Acknowledgements

We thank Satish Nair, Nicholas Wu, Ido Golding, Raven Huang, Wilfred van der Donk, Nick Martin, Andrian Gutu, Lisa Wiltbank for helpful discussions and comments on the manuscript. We would also like to thank Noah Miller for helpful discussions and comments regarding MATLAB. This work was supported by start-up funds from the University of Illinois at Urbana-Champaign and grant K22AI151263 from the National Institutes of Health to J.E.S.

## Contributions

G.C.P., A.M.S, A.S., M.D.K., J.N.R., T.E.K., J.A.I., and J.E.S. designed research. G.C.P., A.M.S, A.S., M.D.K., J.S.P., and J.N.R. performed research. G.C.P., A.M.S, A.S., M.D.K., J.A.I., and J.E.S. analyzed data. G.C.P. and J.E.S. wrote the paper.

## Supplementary Information for

### Materials and Methods

#### Strains, plasmids, and growth conditions

The bacterial strains used in this paper are described in the Supplementary Table 1. Primers are listed in Supplementary Table 2, and plasmids used are described in Supplementary Table 3. *P. aeruginosa* cultures were grown in liquid LB on a roller drum, and on LB plates (1.5% Bacto Agar) at 37 °C. *S. aureus* cultures were grown in liquid LB supplemented with chloramphenicol (10 μg ml^−1^), and on LB plates with chloramphenicol at 37 °C.

#### Generation of *P. aeruginosa* mutants

Gene deletions were generated using the lambda Red recombinase system as previously described^13,47^. The deletion construct was Gibson-assembled from three PCR products. First, approximately 500 bp upstream of the target insertion site was amplified from PA14 genomic DNA. Second, a fragment containing *aacC1* ORF flanked by FRT sites was amplified from pAS03. Third, approximately 500 bp downstream of the target insertion site was amplified from PA14 genomic DNA. The Gibson-assembled product was transformed into PA14 cells expressing the plasmid pUCP18-RedS. The colonies were selected on 30 μg ml^−1^ gentamycin, and then the mutants of interest were counter-selected on 5% sucrose, and pFLP2 was used to flip out the antibiotic resistance gene. pUCP18-RedS and pFLP2 were selected for using 300 μg ml^−1^ carbenicillin. The double and triple mutants were created by subsequent deletions performed similarly.

#### Generation of *S. aureus* strains

USA300 Δ*ahpCF* was generated by amplifying the 5’ and 3’ flanking regions (∼1 kb up- and downstream) of *ahpCF* using the indicated primers (Table S2). 5’ and 3’ fragments were cloned into the pKOR1 knockout vector via site-specific recombination. The deletions were created using allelic replacement, as described previously^48^. USA300 JE2 *katA*:erm was obtained from the Nebraska library^49^. USA300 JE2 *katA*:erm Δ*ahpCF* was generated by transducing the *katA*:erm allele via Φ85 phage from USA300 JE2 *katA*:erm. To create the *ahpCF* reporter construct, the *ahpCF* promoter was cloned into the pAH5 vector^50^ via the indicated primers (Table S2). All constructs were verified by sequencing and all mutant strains were confirmed to be hemolytic by growth on TSA blood agar plates.

#### Bacterial Conditioning of LB Media and H_2_O_2_ treatment

*P. aeruginosa* and *S. aureus* were used to condition media for experiments involving their respective bacteria. Media were conditioned by diluting 50 μL of bacteria from an overnight culture into a 5 mL tube (or scaled up at the same ratio) and allowing it to sit for a defined period at 22 °C. Bacterial cells were then filtered out using a Steriflip sterile filter unit (0.22 μm pore size). To generate LB with defined H_2_O_2_ concentrations, LB was first conditioned, and then defined concentrations of H_2_O_2_ were added. For experiments using purified catalase, 8 mg/ml of bovine liver catalase (Sigma) was used.

#### Fabrication of microfluidic devices

Microfluidic devices were created and fabricated using soft lithography techniques. Devices were designed on Illustrator (Adobe Creative Suite) and masks were subsequently printed by CAD/Art Services. Molds were produced on 100mm silicon wafers (University Wafer) and then spin coated with SU-8 3050 photoresist (MicroChem). Polydimethylsiloxane (PDMS) chips were plasma-treated for bonding to glass slides at least 24 hours before experiments. The devices used in all other experiments discussed above using the *fro* reporter and P_*ahp*_ reporter contained 7 parallel channels 500 μm wide x 50 μm tall x 2 cm long. Long channel experiments used device that were 500 μm wide x 50 μm tall x 27 cm long. Each channel individually contained an inlet and an outlet. These chips were plasma bonded to a 60 mm x 35 mm x 0.16 mm superslip micro cover glass (Ted Pella, Inc.).

#### *P. aeruginosa* in microfluidic devices

All experiments involving the *fro* reporter were conducted using cells at an optical density of approximately 0.5. Cells were injected into the microfluidic device using a pipette and were allowed to settle in the device for 10 minutes prior to exposure to flow. The device set-up involves the use of plastic 5 mL syringes (BD) with attached tubing connecting the needle to the inlet of the device (BD Intramedic Polyethylene Tubing; 0.38 mm inside diameter, 1.09 mm outside diameter). These syringes were situated on a syringe pump (KD Scientific Legato 210) which was used to produce fluid flow. The outlet of the device employed the same tubing and vacated into a bleach-containing waste container. The syringe pump was used to generate flow rates of 0.1-10 μL/min, which correspond to shear rates of 8 – 800 s^−1^.

#### *S. aureus* in microfluidic devices

Experiments measuring P_*ahp*_ expression in *S. aureus* reporter use cells from mid-log phase culture. Flow media was conditioned in *S. aureus* cells for at most 1 hour. Conditioned LB was filtered of cells and supplemented with 8 μM H_2_O_2_ before loading into 5 mL syringe. Cells were injected directly into the flow chamber inlet with a pipette and allowed to settle for 10 min. Flow devices were setup with exact same methods as described in *P. aeruginosa* in microfluidic devices. For no flow conditions, injected cells in flow chambers were given fresh conditioned LB with 8 μM H_2_O_2_ for 2 min.

#### Shear rate calculations

The shear rate experienced in the microfluidic devices was calculated using the equation:

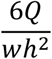

Where Q is flow rate, w is channel width, and h is the channel height.

#### Phase contrast and fluorescence microscopy

Images were obtained on a Nikon Eclipse Ti-2 microscope using the NIS Elements interface. All images were taken with Nikon 40x Plan Apo Ph2 0.95 NA objective, a Hamamatsu Orca-Flash4.0LT camera, and Lumencor Sola Light Engine LED light source.

#### Quantification of *fro*/*ahpCF* expression

The image analysis pipeline employs readily available quantification program (OUFTI) as well as novel code written in MATLAB (Mathworks). Using OUFTI, cell meshes were developed and then used to quantify fluorescence on the YFP and mCherry (or solely YFP in the case of the *S. aureus* experiments). These computed values were then extracted and averaged in MATLAB to yield the per-frame average fluorescence intensity of all cells meshed (>100 per frame) across 3 technical replicates. The ratio of YFP/mCherry or YFP/YFP were then taken to obtain a representative induction value of *P. aeruginosa fro* expression and *S. aureus* P_*ahp*_ expression.

#### Mathematical simulations

To simulate the advection-diffusion of H_2_O_2_ molecules in the microfluidic channel, we combined the laminar transport due to flow with a Brownian dynamics simulation to capture the diffusive behavior. Initially, the simulated channel was seeded with molecules at random positions according to the concentration *c*. Flow was then modeled with a parabolic flow speed profile 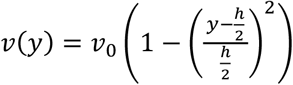 according to the Hagen-Poiseuille equation, with the channel height *h* and width *w* and maximum velocity 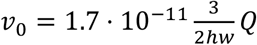 in the center of the channel. In time steps of Δ*t* = 1 ms, each particle was then displaced along the channel according to its lateral position *v*(*y*). In addition, each particle was allowed to diffuse the distance 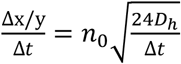 lateral (y) and along (x) the channel, where *n*_0_ is a random number drawn from a uniform distribution in the interval −0.5 ≤ *n*_0_ ≤ 0.5 and 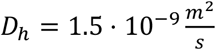 is the diffusion coefficient of H_2_O_2_.

For the simulations, we estimated that 1 of every 100 molecules that reached the channel bottom (where cells are located) were removed. Based on the simulated results, we confirmed that our estimate was reasonable based on the following logic: Figure 3 shows that cells at OD of ∼0.5 remove a majority of H_2_O_2_ in 30 seconds, our calculations show that cells in microfluidic channels are also at an OD of ∼0.5, our calculations show that media flowing at 80 sec^−1^ resides in the channel for ∼30 seconds, and our simulation demonstrates that a shear rate of 80 sec^−1^ results in scavenging of a majority of H_2_O_2_ molecules using our 1 in 100 estimate.

**Figure S1:**
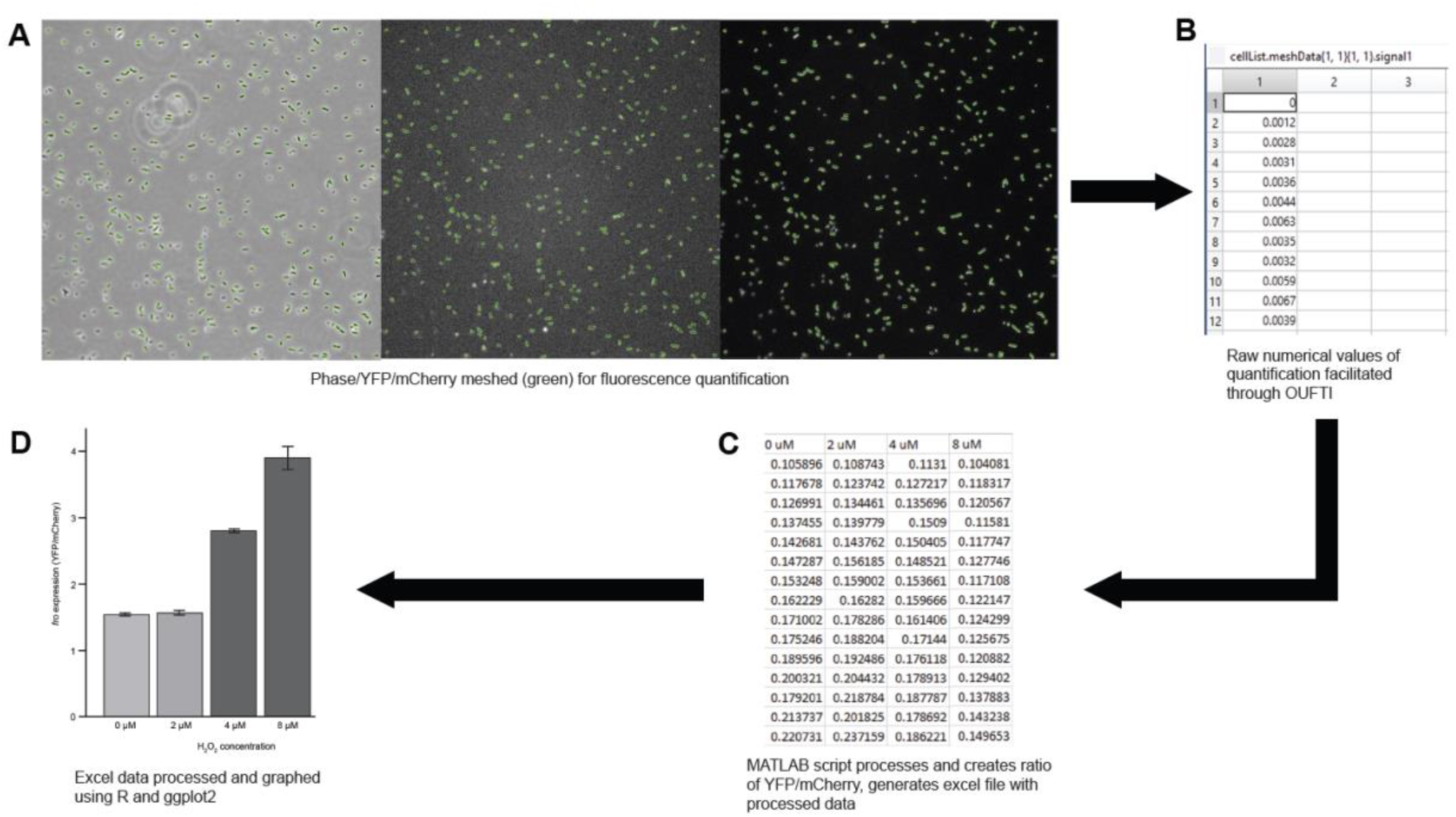
Computational workflow for fluorescence based single cell microscopy quantification. **(A)** Raw microscopy images demonstrating meshes generated through OUFTI from phase image. Meshes overlayed over YFP and mCherry fluorescent images. **(B)** Raw MATLAB output of quantified fluorescent values post-processing with OUFTI. **(C)** Data post-processing with in-lab MATLAB code to yield final values. **(D)** Representative plot generated using data yielded from **C**.

**Figure S2:**
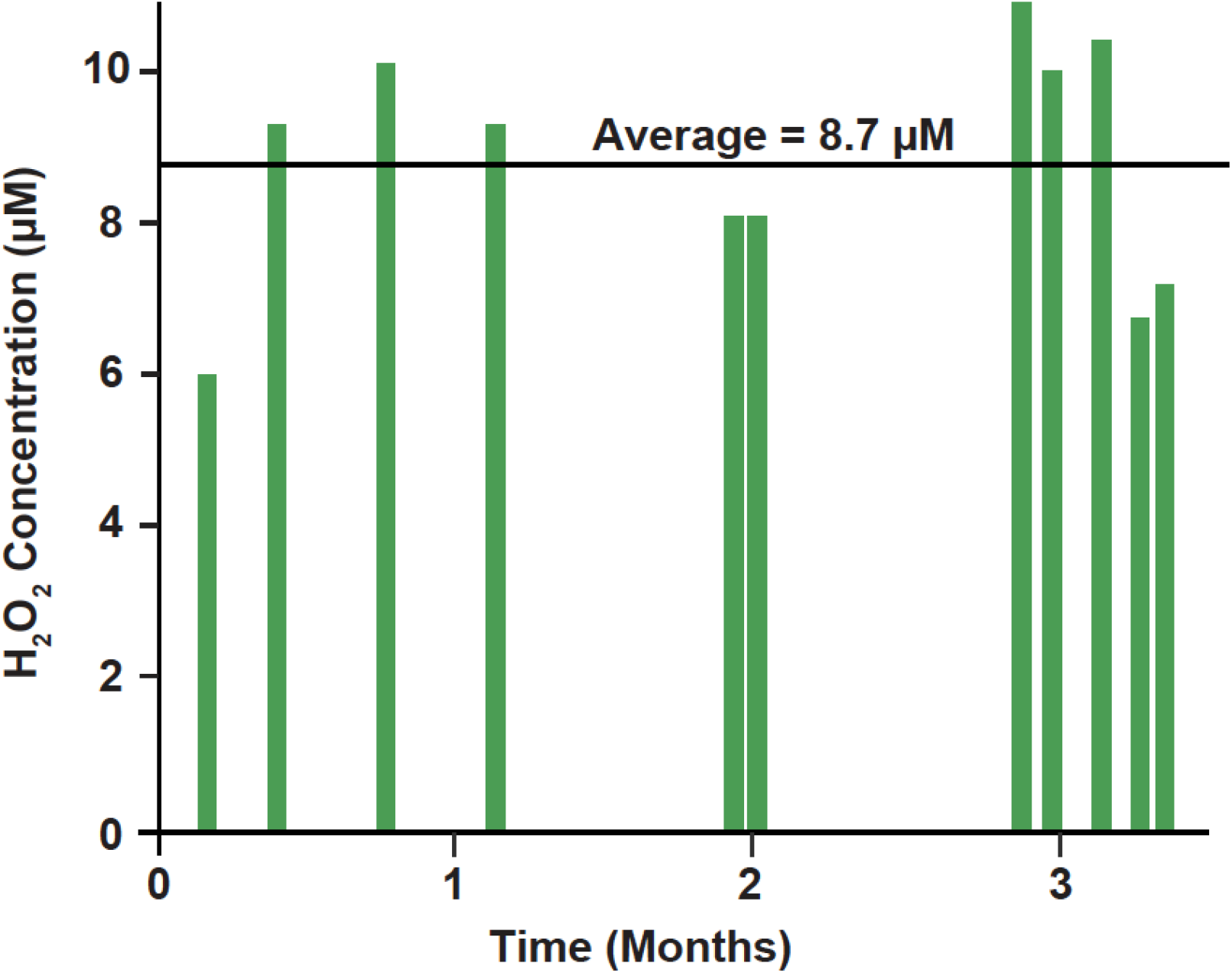
LB contains variable amounts of H_2_O_2_ in standard laboratory conditions. H_2_O_2_ concentration of LB from laboratory storage measured over a period of months. H_2_O_2_ concentrations were measured using a peroxidase assay^21^.

**Figure S3:**
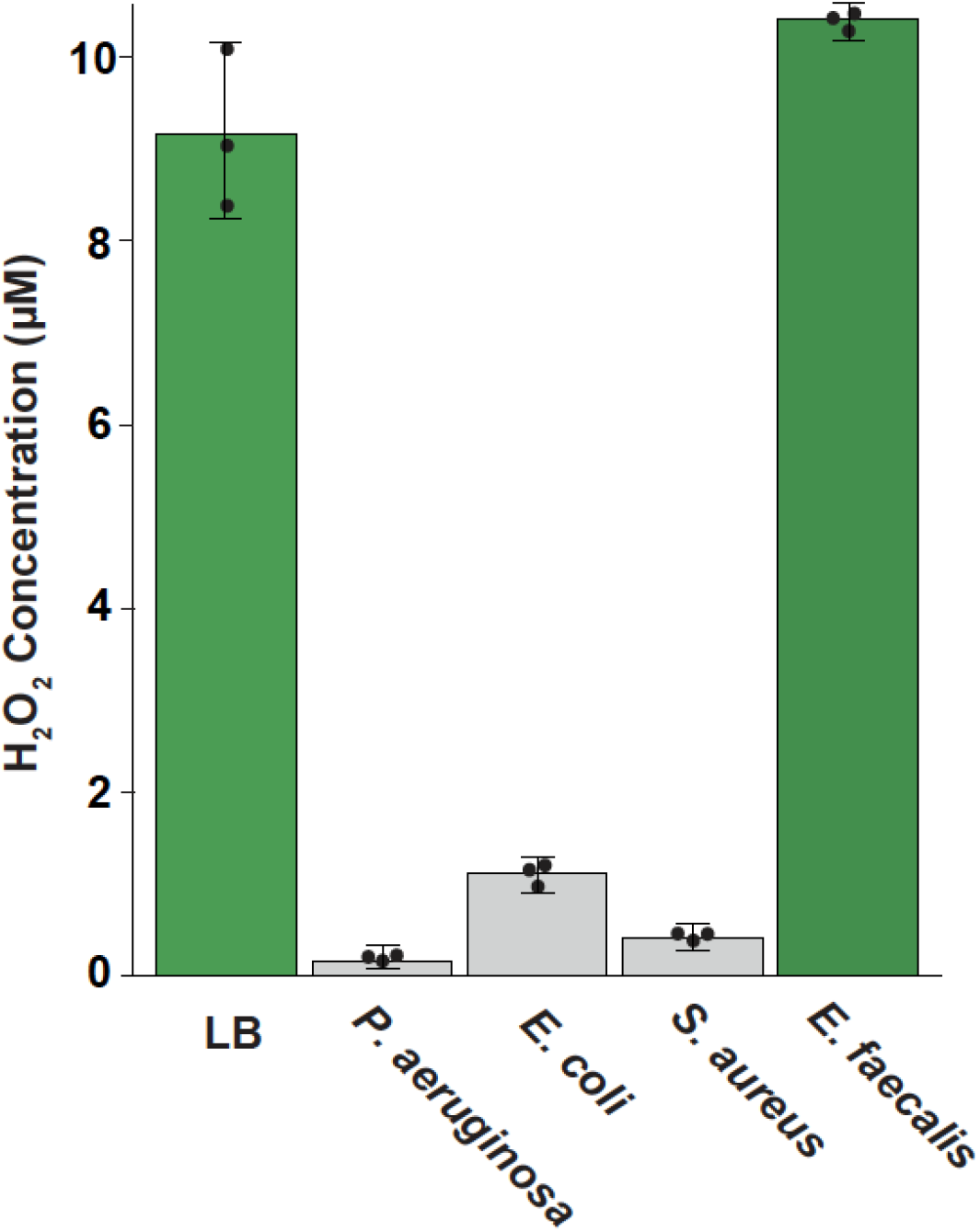
Bacteria scavenging of H_2_O_2_ is conserved, but not universal. H_2_O_2_ concentration of LB when untreated, or treated with ∼0.5 OD wild-type *P. aeruginosa, E. coli, S. aureus*, or *E. faecalis* for 30 minutes. H_2_O_2_ concentrations were measured using a peroxidase assay^21^. Quantification shows the average and standard deviation of three biological replicates.

**Figure S4:**
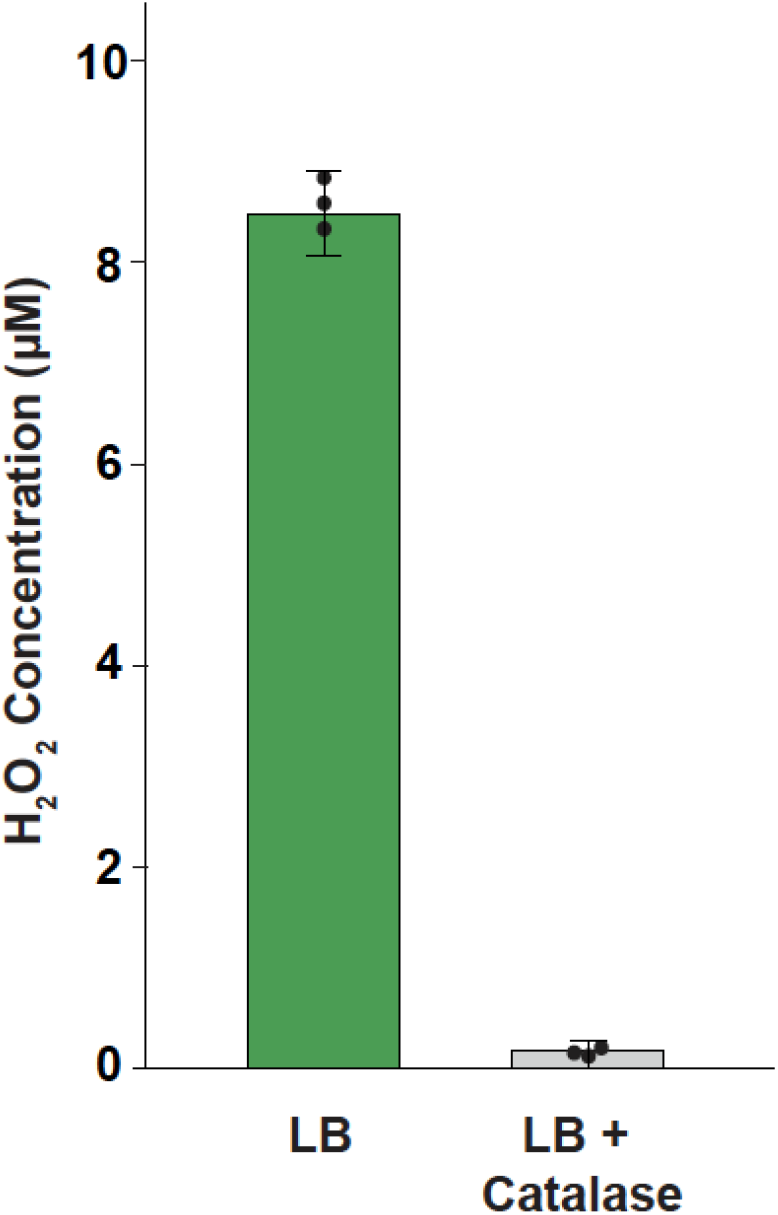
Addition of purified catalase to LB media depletes H_2_O_2_. H_2_O_2_ concentration of LB when untreated or treated with purified catalase. H_2_O_2_ concentrations were measured using a peroxidase assay^21^. Quantification shows the average and standard deviation of three biological replicates.

**Figure S5:**
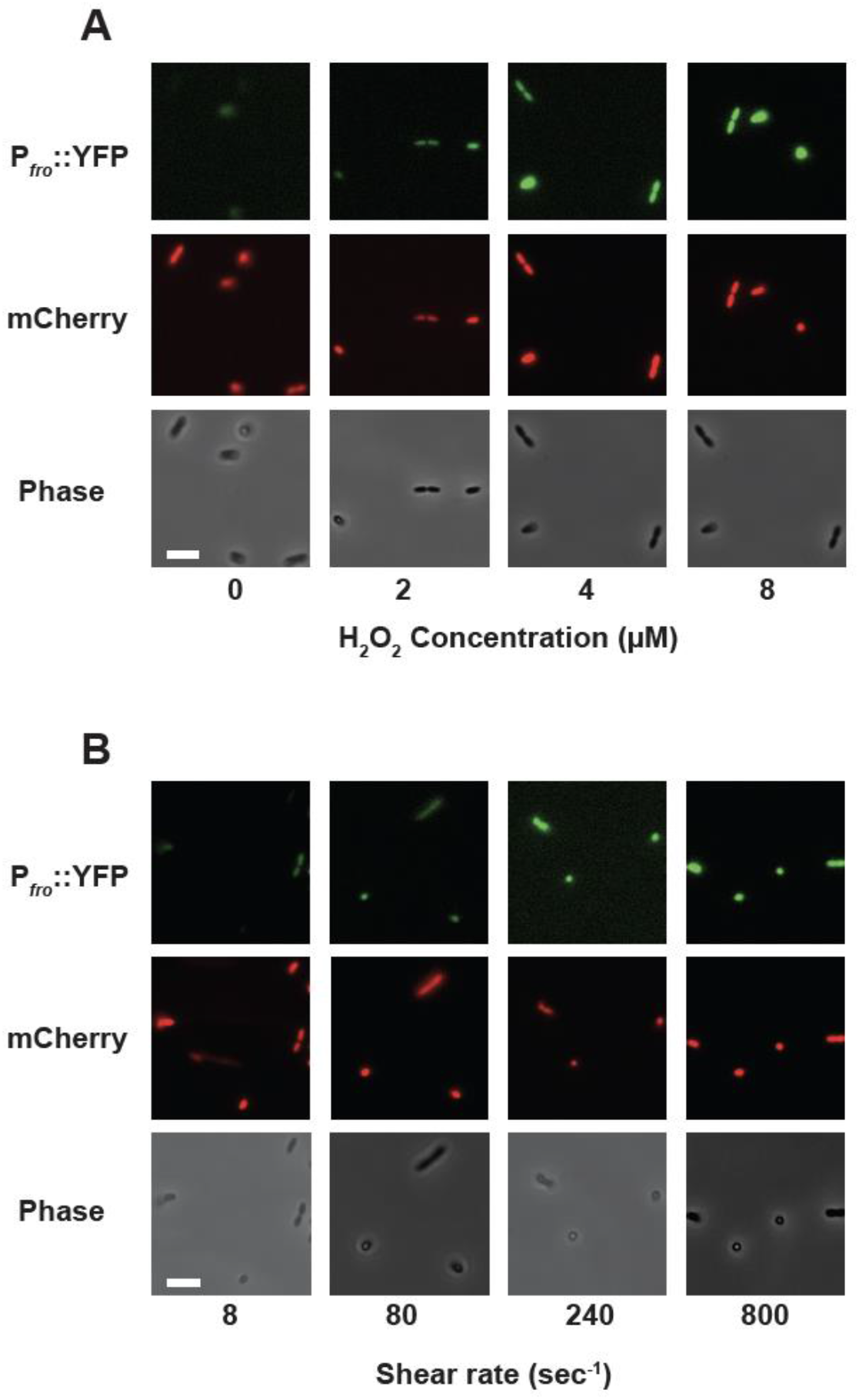
H_2_O_2_ concentration and shear rate tune the flow-sensitive gene expression. **(A)** YFP fluorescence, mCherry fluorescence, and phase contrast images representative of three biological replicates showing *P. aeruginosa* cells treated with 800 sec^−1^ flow and LB with varied H_2_O_2_ concentrations. **(B)** YFP fluorescence, mCherry fluorescence, and phase contrast images representative of three biological replicates showing *P. aeruginosa* cells treated with 8 μM H_2_O_2_ and varied shear rates. Scale bars, 5 μm. Channels are 50 μm tall x 500 μm wide.

**Figure S6:**
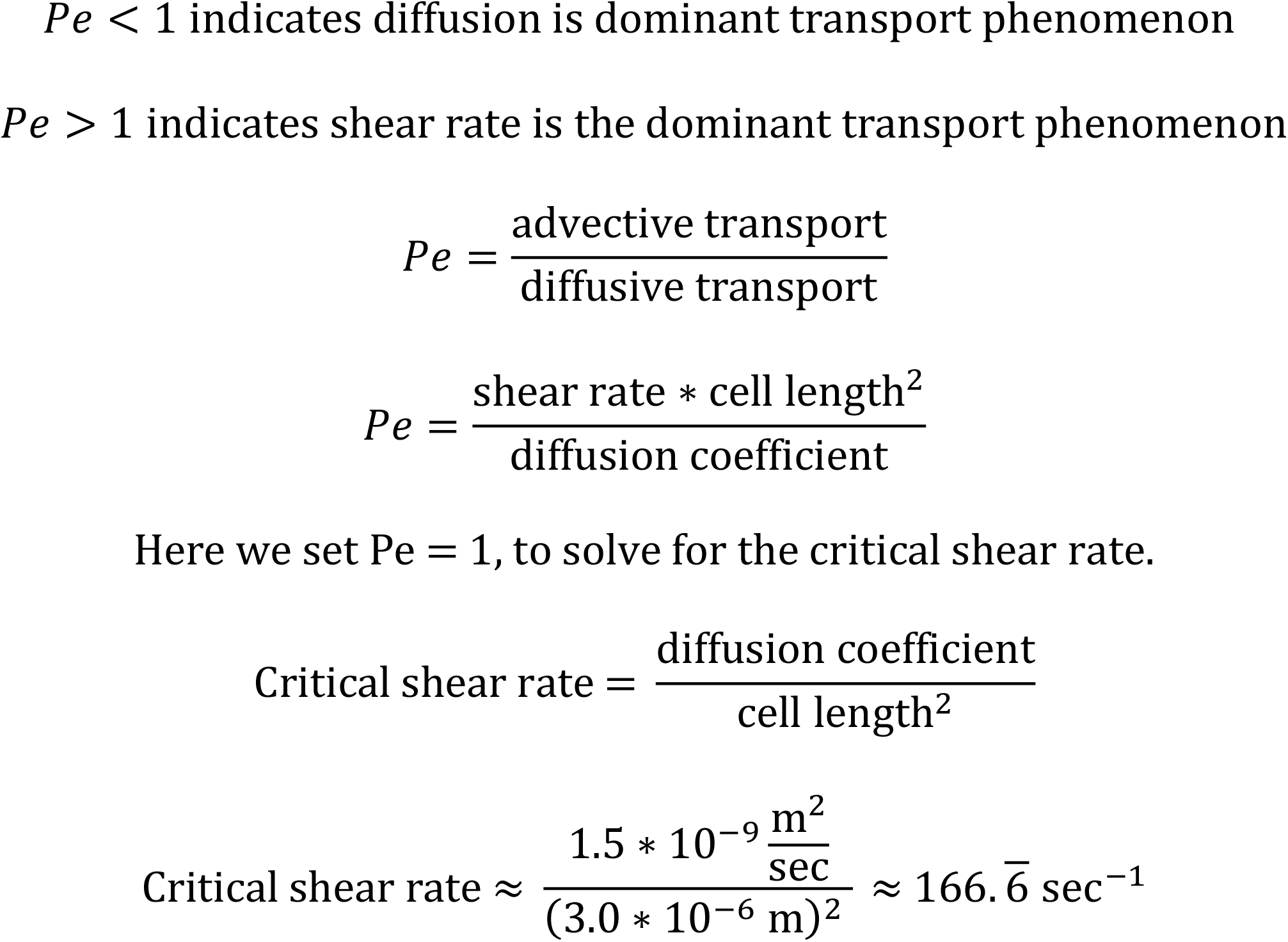
Péclet number calculation. Critical shear rate was calculated by setting the Péclet number = 1. The diffusion coefficient of H_2_O_2_ was estimated to be 1.5 × 10^−9^ m^2^/sec^38^. The cell length of *P. aeruginosa* was estimated to be 3 μm. Cell length is used a characteristic length that defines the relevant scale of the system.

**Figure S7:**
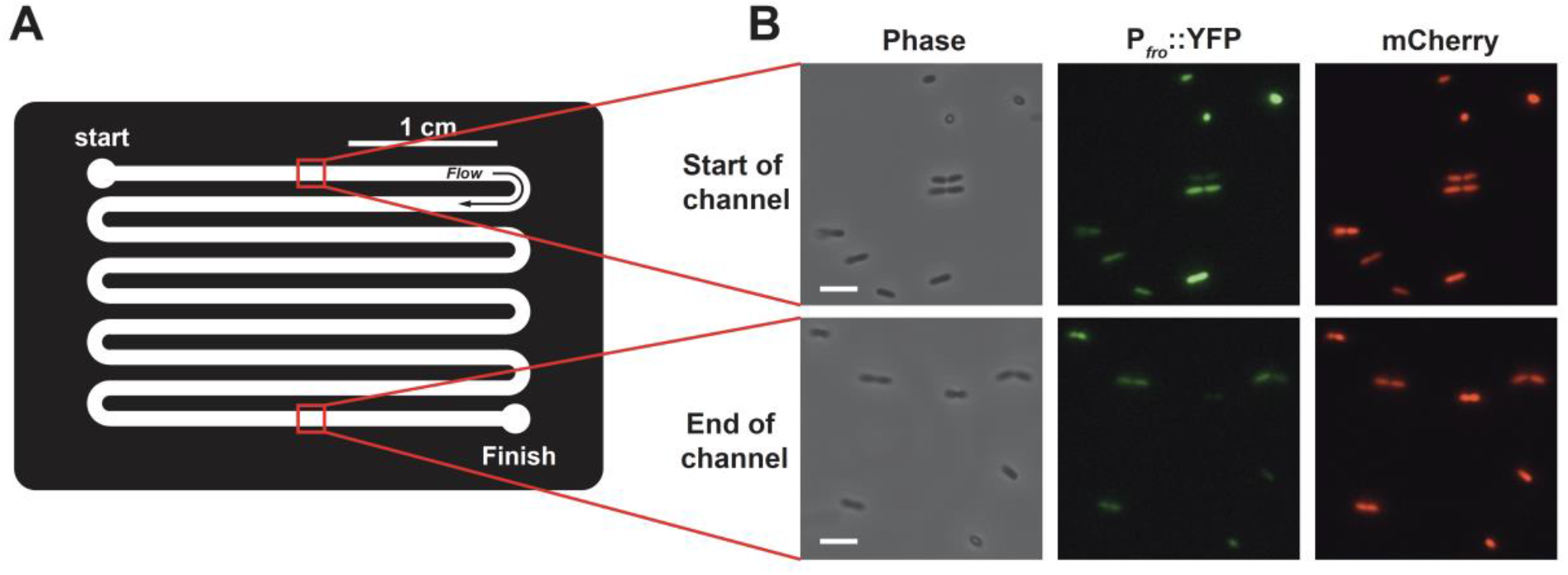
Flow generates gradients of H_2_O_2_ and gene expression. **(A)** Schematic showing top-down view of the long channel microfluidic device used in Figure 4E and in panel B. **(B)** Phase contrast, YFP fluorescence, and mCherry fluorescence images representative of three biological replicates showing *P. aeruginosa* cells treated with 240 sec^−1^ flow and LB with a H_2_O_2_ concentration of 8 μM. Images are from lane 1 (start of channel) and lane 9 (end of channel). Scale bars, 5 μm. Channels are 50 μm tall x 500 μm wide x 27 cm long.

**Figure S8:**
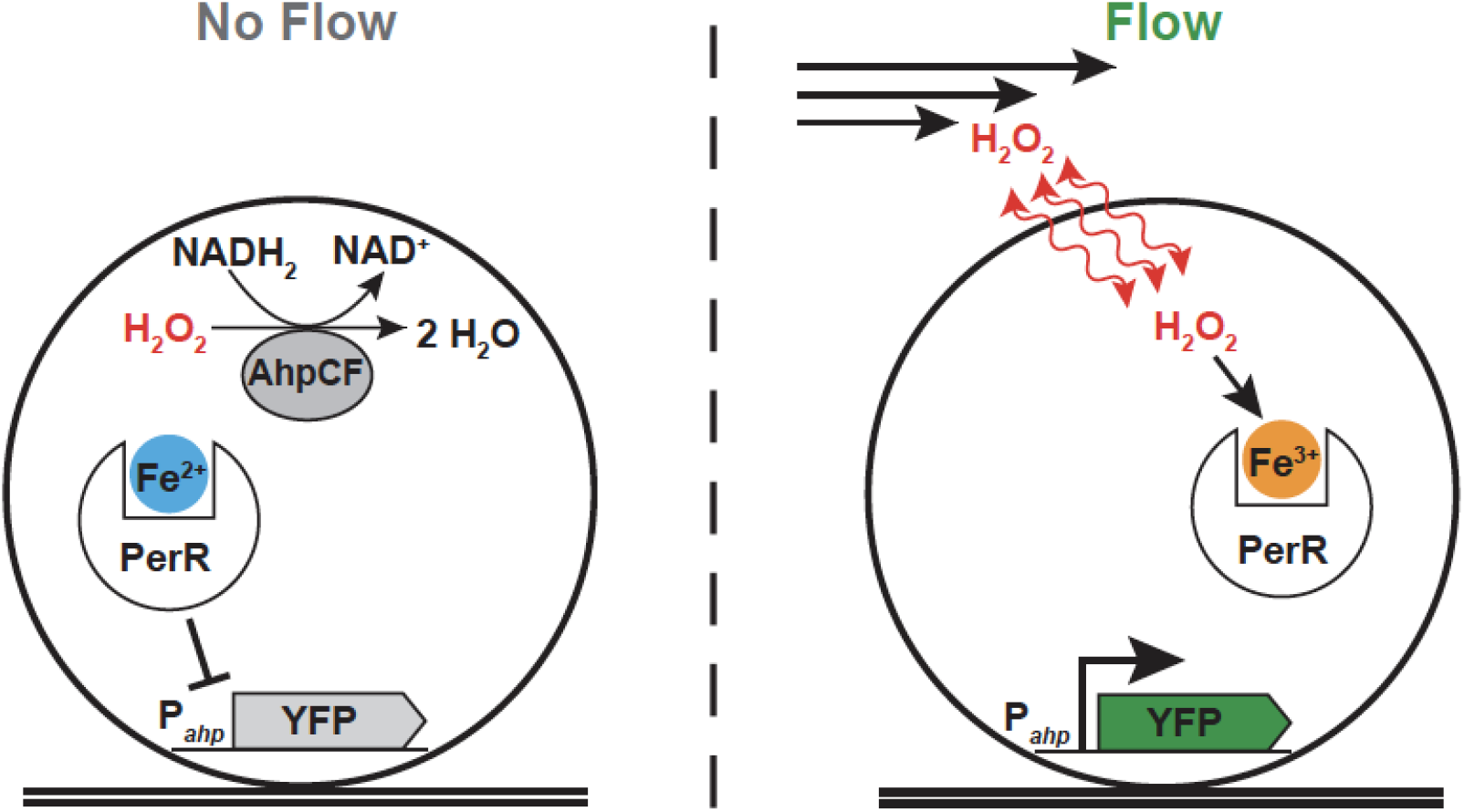
Regulatory model for flow-sensitive gene expression in *S. aureus*. In conditions without flow, H_2_O_2_ is removed from the intracellular environment by the enzymes KatA (not shown here) and AhpCF. With low intracellular H_2_O_2_, the transcriptional regulator PerR is bound to Fe^2+^ and represses transcription from the *ahp* promoter. In conditions with flow, H_2_O_2_ is replenished in the extracellular environment, which leads to a diffusion-driven accumulation of H_2_O_2_ in the cell. With high intracellular H_2_O_2_, Fe^2+^ is oxidized to Fe^3+^. When the transcriptional regulator PerR is bound to Fe^3+^, the *ahp* promoter is derepressed. Thus, flow leads to an induction of *ahpCF* transcription, which we observe as an increase in cytoplasmic YFP.

**Table S1:** Strains used in this study.

| Strain | Description | Source |
| --- | --- | --- |
| ***P. aeruginosa*** |  |  |
| PA14 | wild-type; clinical isolate from burn wound | 51 |
| JS16 | *froA::YFP FRT attB::[plac-mCherry FRT]* | 13 |
| JS230 | $\Delta katB::aacC1$ | This paper |
| JS231 | $\Delta katB::FRT$ | This paper |
| JS232 | $\Delta ahpCF::aacC1$ | This paper |
| JS233 | $\Delta ahpCF::FRT$ | This paper |
| JS234 | $\Delta ahpCF::FRT$ pUCP18-RedS | This paper |
| JS235 | $\Delta ahpCF::FRT \Delta katB::aacC1$ | This paper |
| JS236 | $\Delta ahpCF::FRT \Delta katB::FRT$ | This paper |
| JS253 | $\Delta katA::FRT$ | This paper |
| JS254 | $\Delta katB::FRT \Delta katA::FRT$ | This paper |
| JS255 | $\Delta ahpCF::FRT \Delta katA::FRT$ | This paper |
| JS256 | $\Delta ahpCF::FRT \Delta katB::FRT \Delta katA::FRT$ | This paper |
| ***S. aureus*** |  |  |
| USA300 JE2 | wild-type; used for peroxide conditioned media experiments. | 49 |
| JS244 | $\Delta ahp$ | This paper |
| JS251 | *katA::erm* | This paper |
| JS252 | $\Delta ahp, katA::erm$ | This paper |
| JS258 | pAH5:*ahp* | This paper |
| ***E. coli*** |  |  |
| MG1655 | wild-type; used for peroxide scavenging experiments | *E. coli* Genetic Stock Center |
| AG75 | Hpx- $\Delta ccp$; $\Delta ahpF::kan$, $\Delta(katG17::Tn10)1$, $\Delta(katE12::Tn10)1$, $\Delta ccp$ | 31 |
| ***E. faecalis*** |  |  |
| OG1RF | wild-type; used for peroxide conditioned media experiments. | ATCC 47077 |

**Table S2:**
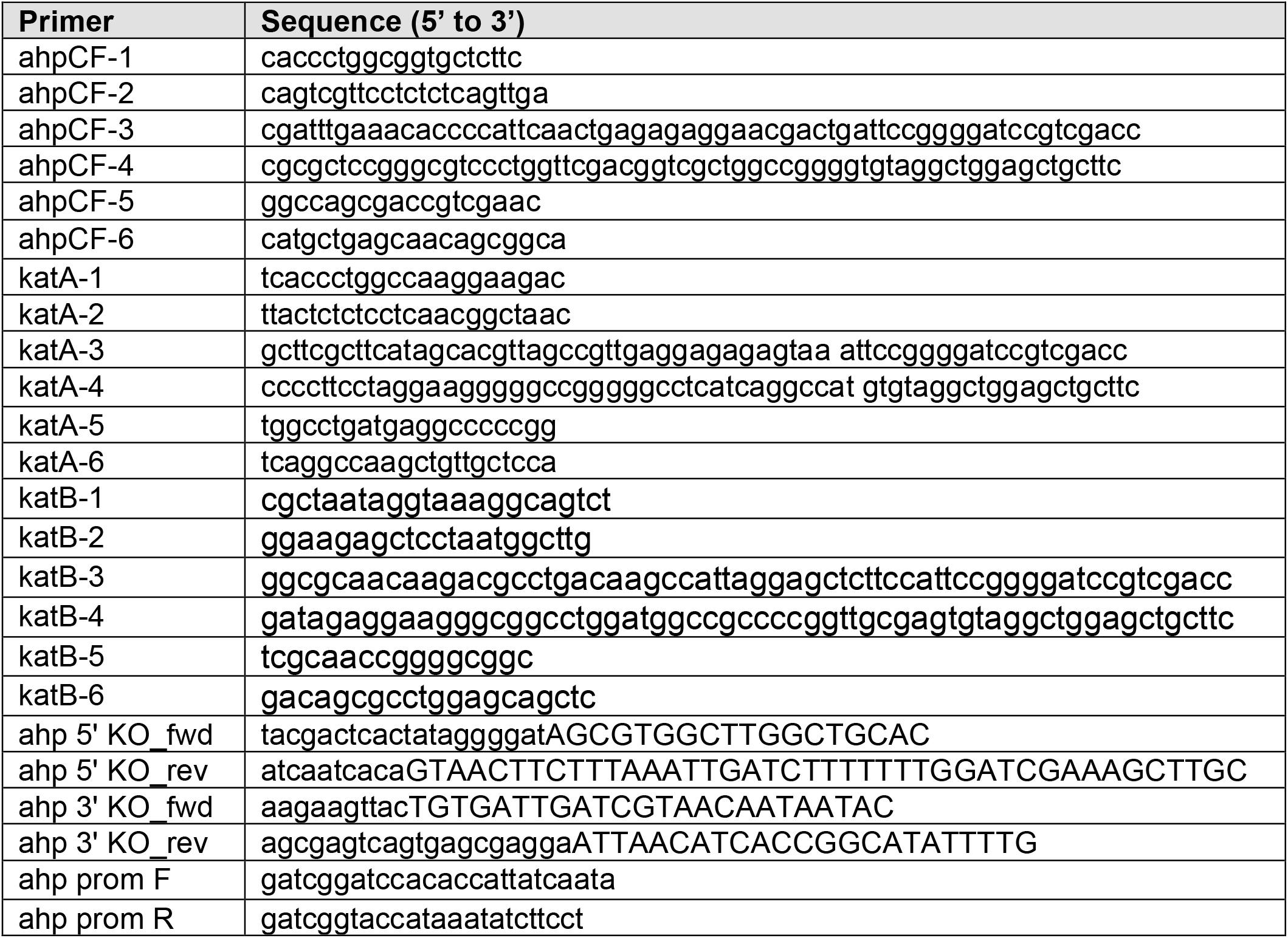
Primers used in this study.

**Table S3:**
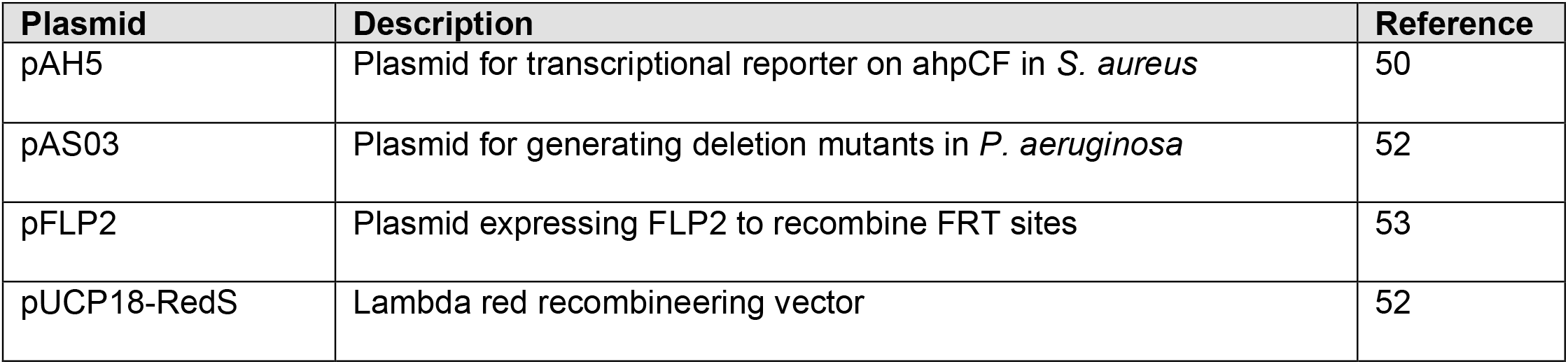
Plasmids used in this study.

